## Supplementary Material for "SNAI2 cooperates with MEK1/2 and HDACs to suppress BIM- and BMF-dependent apoptosis in *TERT* promoter mutant cancers"

Begränsad delning

SUPPLEMENTAL DATA

### Suppression of SNAI2 synergizes with inhibitors of MEK1/2 and HDACs to sensitize *TERT* promoter mutant cancers to BIM- and BMF-dependent apoptosis

A. Tandon and J.L. Stern

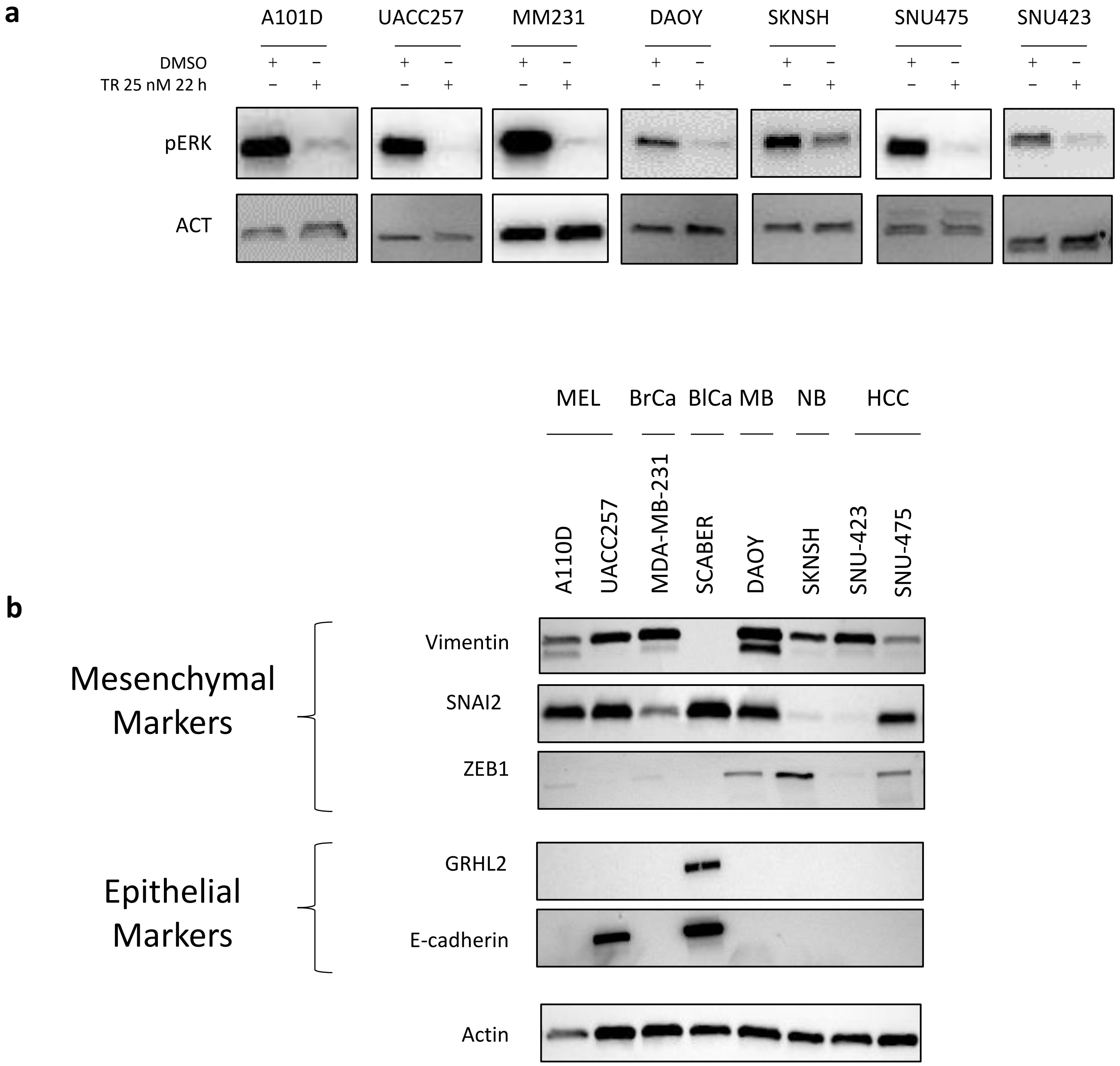

Fig S1. **ERK activation in TPM Cancer Cells is Inhibited by Low Doses of MEK1/2 inhibition (MEKi).** (a) MEK1/2 was inhibited in cells by treatment with trametinib (TR) 25 nM for 22 hours followed by immunoblot for phospho-ERK (pERK). (b) Mesenchymal (Vimentin, Slug and Zeb1) and epithelial (GRHL2 and E-cadherin) markers were tested in different cell lines by western blots. A10D and UACC257 (Melanoma), MDA-MB-231 (Breast cancer), SCaBER (Bladder cancer), DAOY (Medulloblastoma), SKNSH (Neuroblastoma), SNU423 and SNU475 (Hepatocellular carcinoma). β-Actin was used as loading control for Western blots. MEL – Melanoma; HCC – Hepatocellular carcinoma; BrCa – Breast cancer; BlCa – Bladder cancer; MB – Medulloblastoma; NB – Neuroblastoma.

**a**

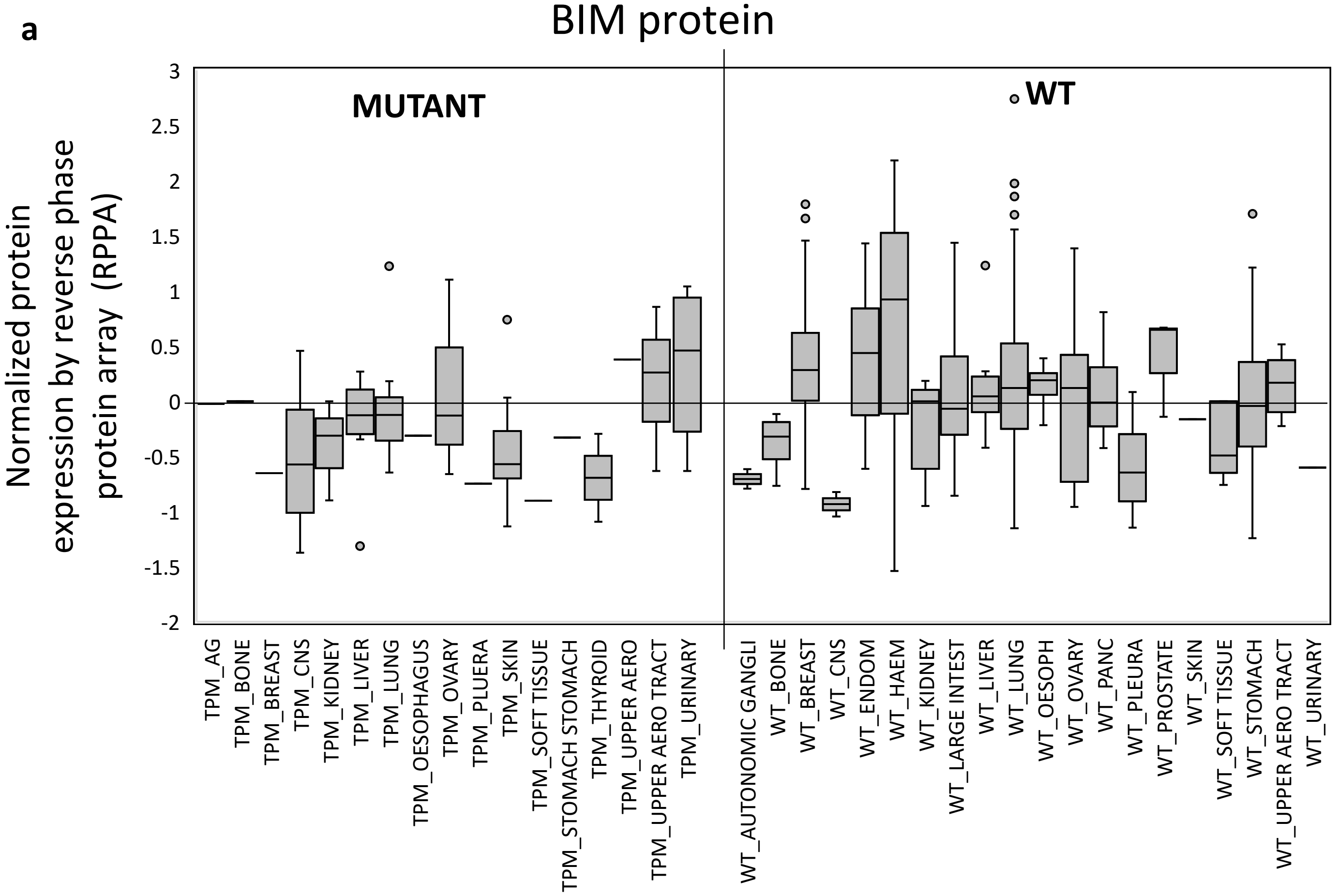

**b**

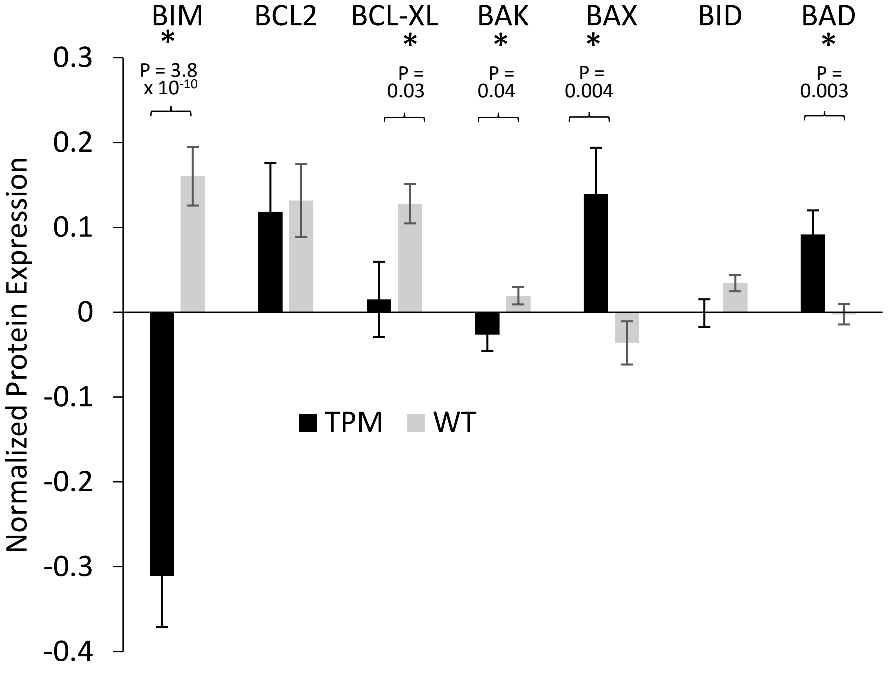

Fig S2. **BIM protein is below average in most TPM cancer types.** (a) Reverse phase protein array (RPPA) data was obtained from Cancer cell line encyclopedia (CCLE). Cell lines annotated as *TERT* promoter mutant or wild type (WT) were analyzed for BIM (*BCL2L11*) protein expression. Graph shows normalized data. (b) Data from RPPA depicting expression of BIM and other Bcl2-family member proteins in *TERT* promoter mutant (TPM, black) and wild type (WT, grey) cell lines.

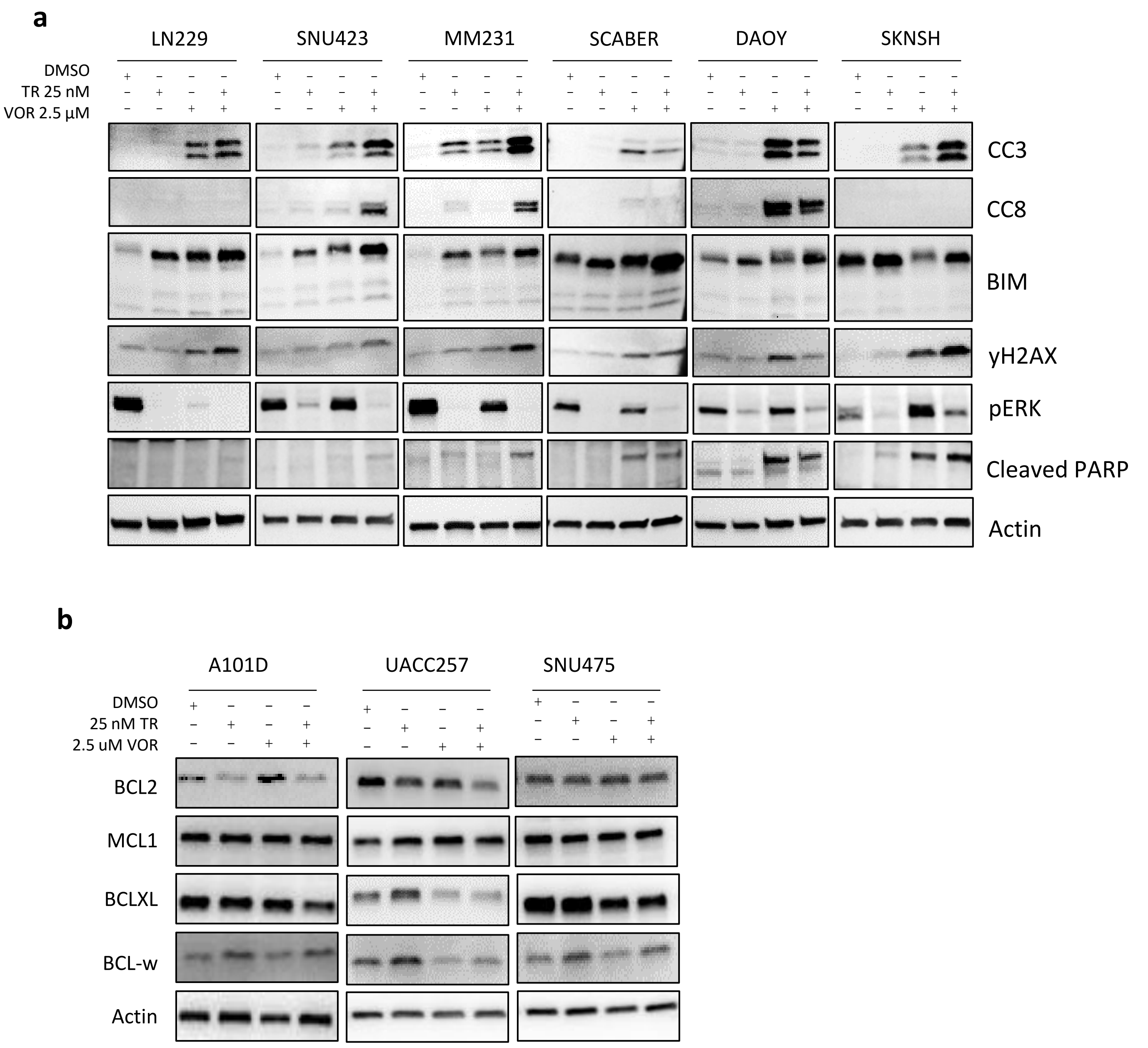

Fig. S3 **Combining MEKi and HDACi induces apoptosis in TPM cells.** (a) Cells were treated for 24 hours with trametinib (TR, 25 nM) with or without vorinostat (VOR, 2.5 µM) and probed for apoptosis (CC3) and induction of DNA damage (pH2AX) on Western blots. Blots were also probed for cleaved caspase-8 (CC8), phospho-ERK (pERK), BIM and cleaved PARP. (b) Cell lines treated in (a) were probed by immunoblots for pro-survival markers Bcl2, MCL1, BCL-XL and BCL-W. β-Actin was used as a loading control and for normalization.

### Table S1.

| GENE | A101D | UACC257 | SNU-475 | SNU-423 | MDA-MB-231 | DAOY | SK-N-SH | SCaBER |
|---|---|---|---|---|---|---|---|---|
| ***TERT*** | c.228C>T (-124C>T)* in promoter PMID: 31068700 | c.250C>T (-146C>T)* in promoter PMID: 31068700 | c.228C>T (-124C>T)* in promoter PMID: 31068700 | c.228C>T (-124C>T)* in promoter PMID: 31068700 | c.228C>T (-124C>T)* in promoter PMID: 31068700 | c.228C>T (-124C>T)* in promoter PMID: 31068700 | c.228C>T (-124C>T)* in promoter PMID: 26171145 | c.228C>T (-124C>T)* in promoter PMID: 24035680 PMID: 31068700 |
| **RAS pathway** | Heterozygous for BRAF | Heterozygous for BRAF | | | Heterozygous for BRAF Heterozygous for KRAS | | | |
| ***MGMT*** | MUT PMID: 30709805 | WT PMID: 30709805 | | | | | | |
| **TP53** | | Has no TP53 CCLE; Cosmic-CLP | MUT p.Asn239Asp (c.715A>G), p.Cys275Arg (c.823T>C) and p.Asn288Ser (c.863A>G) PMID: 8824565 | MUT c.376-2A>G; splice acceptor mutation (PMID: 8824565; CCLE | Homozygous MUT p.Arg280Lys (c.839G>A) PMID: 15900046 PMID:16541312 PMID:17088437PMID: 18277095 PMID:28889351 ATCC | | | Homozygous MUT p.Arg110Leu (c.329G>T) PMID: 850064; CCLE; Cosmic-CLP |
| ***CDKN2A*** | | CDKN2A deletion PMID: 29492214 | | | Homozygous for CDKN2A deletion PMID: 19593635 | Homozygous for CDKN2A deletion (ATCC). | | |
| ***ALK*** | | | | | | | p.Phe1174Leu (c.3522C>A) ClinVar=VCV000217852) PMI: 18724359; PMID: 28350380. | |

*-124 or -146 refers to base positions upstream of the *TERT* ATG. c228 of c250 refer to nucleotide positions in HG38 chromosome, 5: base 1,295,228 or base 1,295,250.

Table S1. **Cell lines used in this study.** Data from Cellosauraus (https://www.cellosaurus.org/) showing the mutational signature of different cell lines used in the study.
